# Akkermansia muciniphila Impacts Group B Streptococcus Vaginal Colonization

**DOI:** 10.1101/2025.09.18.677025

**Authors:** Stephanie M. Marroquin, Shirli Cohen, Melody N. Neely, Kelly S. Doran

**Affiliations:** Department of Immunology and Microbiology, University of Colorado School of Medicine, Aurora, CO, USA; Department of Molecular and Biomedical Sciences, The University of Maine, Orono, ME, USA

**Keywords:** Vaginal colonization, Group B *Streptococcus*, *Akkermansia muciniphila*, RNA-seq: Capsular polysaccharide, Probiotic

## Abstract

*Streptococcus agalactiae* or Group B *Streptococcus* (GBS) is an opportunistic pathogen that asymptomatically colonizes the vaginal tract of up to 30% of healthy individuals. However, during pregnancy it is associated with adverse pregnancy outcomes, and GBS can be transmitted to the fetus *in utero*, or the newborn during vaginal birth, resulting in invasive neonatal disease. Previously we identified that *Akkermansia muciniphila* alters the dynamics of GBS vaginal colonization. However, the global effect of *A. muciniphila* in the female genital tract remains largely unknown, as it has predominantly been studied in the context of gastrointestinal health. Here we determine that *A. muciniphila* promotes GBS aggregation and attachment to human vaginal epithelial cells (hVECs). RNA-sequencing analysis revealed that *A. muciniphila* changed expression of 258 unique GBS genes during hVEC colonization, with many involved in cell wall/membrane/envelope biogenesis. We demonstrate that *A. muciniphila*-mediated increases in GBS aggregation and attachment to hVECs are dependent on GBS capsule and pili, respectively. Lastly, we found that *A. muciniphila* promoted GBS aggregation in the murine vaginal lumen and continual treatment with *A. muciniphila* reduced GBS vaginal persistence. Our results highlight the dynamic impact of *A. muciniphila* on GBS gene expression and vaginal colonization, as well as demonstrate the probiotic potential of *A. muciniphila* in the vaginal environment.

**IMPORTANCE:** Group B *Streptococcus* (GBS) is a frequent colonizer of the vaginal tract of healthy people, however during pregnancy, maternal colonization is associated with adverse pregnancy outcomes. GBS is a leading cause of neonatal sepsis and meningitis and can be transmitted to a newborn during vaginal delivery or by ascension into the uterus during pregnancy. Influence of the vaginal microbiota on GBS pathogenesis remains greatly underappreciated. We have found that GBS is associated with the mucin-degrading intestinal commensal *Akkermansia muciniphila*, a newly identified colonizer of the vaginal tract. Significance of our research is in identifying the mechanistic impact of this commensal organism on GBS aggregation, cell adherence, and gene expression, as well as its therapeutic potential during GBS vaginal colonization. Unraveling relationships between GBS and the vaginal microbiota will improve maternal and fetal health and may enterprise the development of alternative methods to reduce GBS *in utero* complications and neonatal disease.

## INTRODUCTION

*Streptococcus agalactiae* or Group B *streptococcus* (GBS) is a Gram-positive, β-hemolytic, opportunistic pathogen that asymptomatically colonizes the female genital tract (FGT) and gastrointestinal (GI) tract of 25-30% of healthy women ^1^. During pregnancy GBS is associated with adverse pregnancy outcomes including preterm premature rupture of membranes (PROM), chorioamnionitis, stillbirth, and preterm birth ^2–4^. Notably, the majority of preterm births are due to ascending microbial infections, with 10% of these being caused by GBS ^5^. Maternal GBS GI and/or vaginal tract colonization is the primary risk factor in neonatal GBS disease, and approximately 50% of GBS colonized mothers deliver newborns that are also colonized with GBS ^6,7^. GBS transmission can lead to neonatal pneumonia, sepsis, and meningitis ^8^. GBS encodes for a myriad of virulence factors, including cell envelope associated factors such as pili, serine-rich repeat (Srr) proteins, and capsular polysaccharides (CPS) which may aid host cell binding as well as evading immune responses ^5,9–13^. One of the most notable GBS virulence factors is the CPS, which is uniquely sialylated and provides GBS with the capacity for “molecular mimicry” within the host ^14^. To date, there have been 10 identified capsular serotypes, with Ia, Ib, II, III, and V being the most associated with disease worldwide^1^. Importantly, a large proportion of neonatal meningitis is caused by serotype III strain ^15,16^.

During pregnancy GBS FGT colonization may be intermittent and transient ^17^. This is likely a consequence of a combination of GBS determinants, the commensal microbiota, and host immune responses ^18^. In the vagina, GBS must overcome various challenges in order to successfully colonize the host, including the physical barrier created by mucins on epithelial cells, competition with commensal vaginal microbiota, and mucosal immunity ^19,20^. Importantly, the vaginal microbiota is composed of numerous taxa that vary greatly based on factors such as geography, race or ethnicity, hormone cycles, and pregnancy; thus, the relationship between GBS and the microbiota is quite complex^21^. Recently, a study from our laboratory identified a notable commensal of the GI tract, *Akkermansia muciniphila,* in the murine vaginal microbiota, which helped to promote GBS colonization ^22^, however the mechanisms underlying these observations are not known. Further, examination of human vaginal samples from a pregnancy cohort identified co-occurrence of GBS and *A. muciniphila* in the vaginal tract suggesting that *A. muciniphila* may impact the human FGT^22^. As such, it is of significant importance to investigate key factors that facilitate GBS survival in the FGT as this is a prerequisite to ascending infection and neonatal disease ^5^.

*A. muciniphila* is a Gram-negative, mucin-degrading, anaerobe belonging to the Verrucomicrobiota family that was isolated from human feces in 2004, which can be found in approximately 90% of healthy individuals ^23,24^. Identified as an intestinal symbiont colonizing the mucosal layer, *A. muciniphila* has been studied for its role in the GI tract and its pilin (Amuc_1100) has been shown to be involved in host immunological homeostasis at the gut mucosa ^25–27^. These studies have driven the popularity of *A. muciniphila* as a probiotic, with numerous private companies now selling varieties of live and pasteurized probiotic *A. muciniphila* ^28,29^. Importantly, previous works have also shown that *A. muciniphila* strains can vary in oxygen sensitivity, with Muc^T^ (our strain of interest) being significantly aerotolerant ^30^. Here we examine the mechanisms of *A. muciniphila* and GBS interactions and impact on vaginal colonization. Our observation suggests cooperative and antagonistic effects of *A. muciniphila* on GBS pathogenesis and highlights the complexity of interactions between commensal organisms and opportunistic pathogens in the vaginal niche.

## RESULTS

### GBS and *A. muciniphila* co-aggregate *in vitro* and exhibit increased attachment to human vaginal epithelial cells (hVECs)

To further examine the presence of *A. muciniphila* in the vaginal microbiota during pregnancy in a human cohort, we examined a publicly available metadata set that included 749 vaginal swabs of pregnant women at time of birth from three Parisian hospitals ^31^. Our analysis showed that 4.1% of the 749 women were colonized with *A. muciniphila*. Of the individuals who were positive for *A. muciniphila*, 87.1% were also positive for GBS. These data indicate a 5 times increased likelihood to be GBS positive when positive for *A. muciniphila* (**Figure 1A**). Moreover, upon examining GBS sequencing reads within these data to estimate GBS abundance, samples positive for *A. muciniphila* had higher abundance of GBS than samples that were negative for *A. muciniphila* (**Figure 1B**). Altogether, these data indicate that there is significant co-occurrence between GBS and *A. muciniphila* in the pregnant women of this cohort.

**Figure 1.**
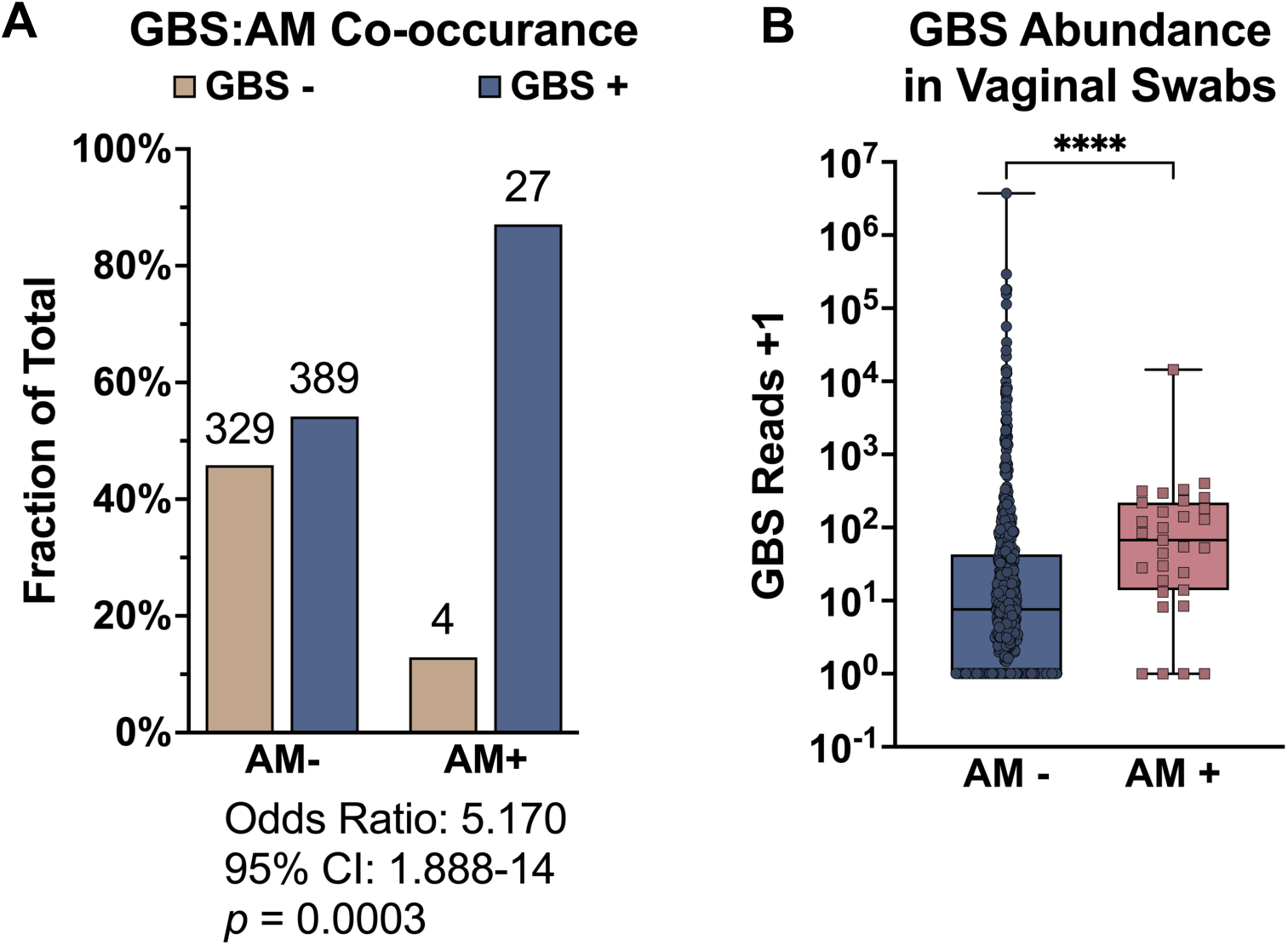
Co-occurrence of GBS and *A. muciniphila* in the vaginal tract of pregnant individuals. **(A)** Co-occurrence of GBS in *A. muciniphila* negative (-) and positive (+) pregnant individuals. Values above the bar indicate the raw number of samples in each group. **(B)** GBS abundance represented by GBS reads in *A. muciniphila* (-) and (+) individuals. Box and whisker plot showing min and max. Statistical analysis was determined by Fisher’s exact test with Baptista-Pike method **(A)** and Mann-Whitney U-test **(B)**. Data points represent study individuals. ****, *p* ≤ .0001.

While these and our previously published results demonstrate that *A. muciniphila* is present in the vaginal tract and influences GBS abundance, whether they are directly interacting is unknown. To begin investigating this, we incubated GBS strain COH1 (serotype III) and *A. muciniphila* in PBS and observed that the two bacteria aggregate together significantly more compared to mono-cultures (**Figure 2A,B,C**). These prominent aggregates can be further visualized by fluorescence microscopy using a GFP-expressing GBS strain and post staining with an antibody against *A. muciniphila* (**Figure 2D**). We observed *A. muciniphila* aggregation with other GBS clinical isolates of varying serotypes (**Figure 3A**) as well as with a cohort of GBS strains isolated from the vaginal tract of pregnant women ^32^ (**Figure 3B**). Interestingly, trends can be observed across the various vaginal isolates we examined based on their serotype, with some serotypes co-aggregating to a slightly higher degree. It is possible that the composition of each capsular serotype plays a role in the propensity to aggregate to *A. muciniphila*, which requires further investigation. We next sought to examine COH1 and *A. muciniphila* interactions on human vaginal epithelial cells using adherence assays ^33^. Here, we found that the presence of *A. muciniphila* significantly increased GBS adherence to hVECs by approximately 1.6 and 2.8-fold for strains COH1 and CJB111 (serotype V) respectively (**Figure 3A,B**). Importantly, we also found that the presence of GBS (COH1) increased *A. muciniphila* adherence to hVECs by approximately 1.8-fold (**Figure 3C**).

**Figure 2.**
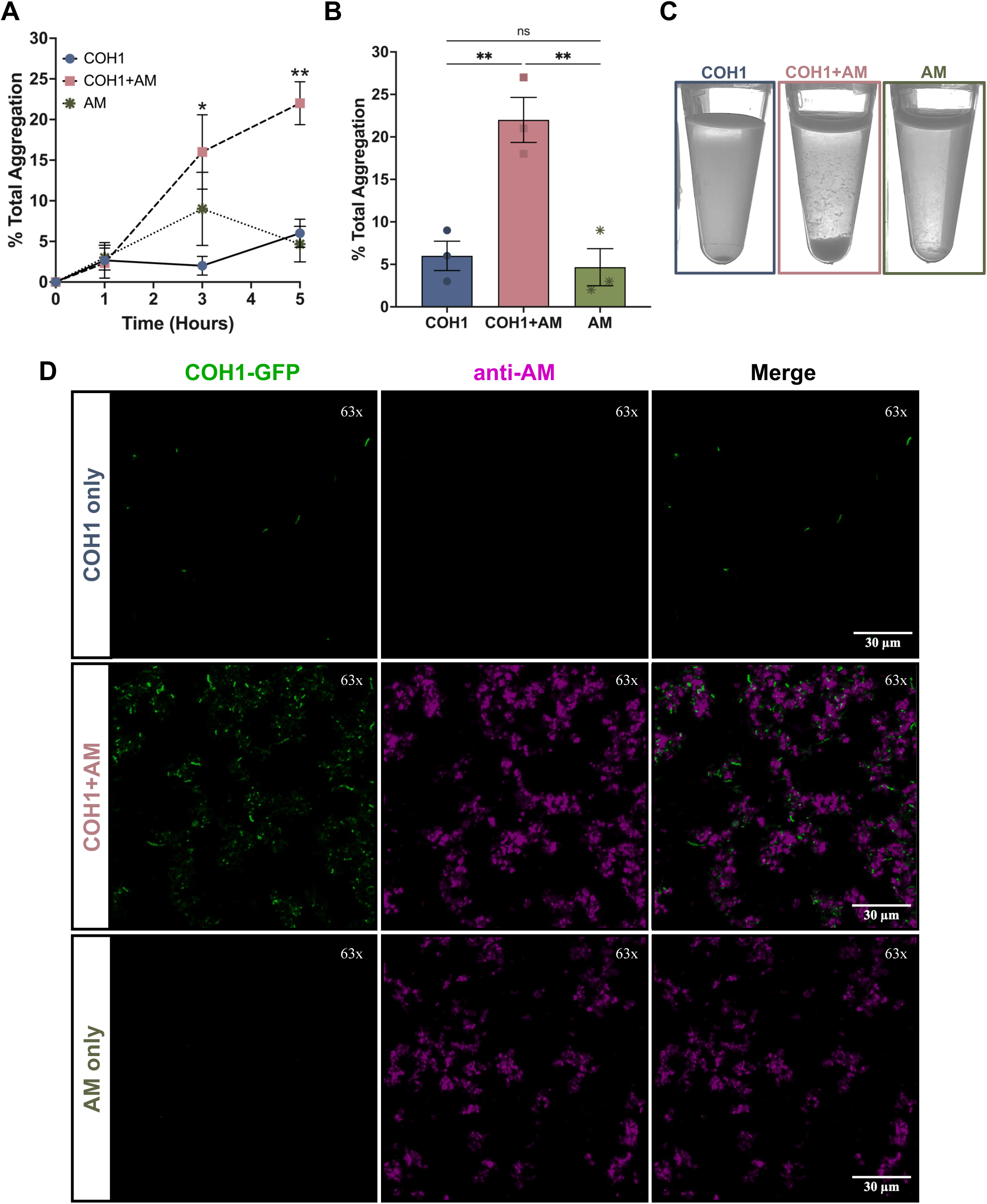
GBS and *A. muciniphila* co-aggregate *in vitro.* **(A)** OD_600_ was monitored at 0h, 1h, 3h, and 5h, and percent aggregation was calculated from initial OD_600_. **(B)** Bar graph and **(C)** image depict aggregation at 5h. **(D)** Aggregates were examined by confocal fluorescence microscopy (representative image). Statistical analysis was determined by Student’s T-test. Data points represent the average of independent experiments. Error bars represent +/-SEM. ns, > .05; *, *p* ≤ .05; **, *p* ≤ .01.

**Figure 3.**
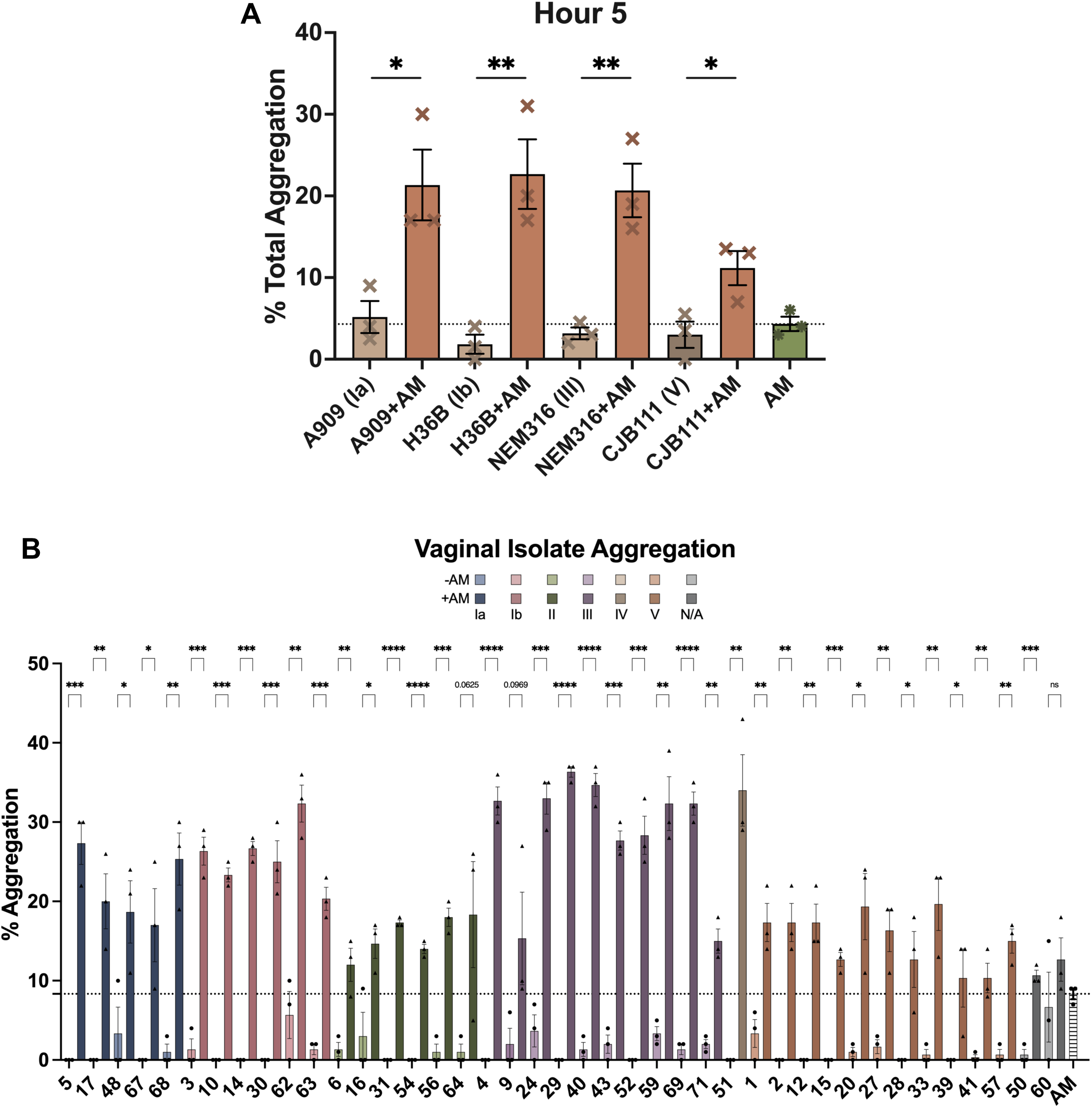
*A. muciniphila* interacts with other GBS capsular serotypes, sequencing types, and GBS vaginal isolates from pregnant individuals. **(A)** Aggregation of additional GBS serotypes and sequencing types to *A. muciniphila* was examined. **(B)** Aggregation of various GBS human vaginal isolates to *A. muciniphila* was assessed. Isolates are color coded based on their capsular serotype. Lighter color and (●) denote the GBS only mono-condition and darker color and (▴) denote the GBS + *A. muciniphila* co-condition. Statistical analysis was determined by Student’s T-test. Data points represent the average of independent experiments **(A)** or technical replicates from independent experiments **(B)**. Error bars represent +/-SEM. ns, > .05; *, *p* ≤ .05; **, *p* ≤ .01; ***, *p* ≤ .001; ***, *p* ≤ .001; ****, p ≤ .0001.

### RNA sequencing reveals unique GBS transcriptome changes during co-infection of hVECs

After observing the ability of GBS and *A. muciniphila* to co-aggregate and bind to hVECs, we next sought to determine the effect of *A. muciniphila* on GBS gene expression during infection of the vaginal epithelium. We performed RNA-sequencing to examine global transcriptomic changes in GBS during mono-infection of hVECs (GBS+hVECs) and co-infection with *A. muciniphila* (GBS+AM+hVECs), (**Figure 5A**) and compared to a GBS only control (GBS grown in Keratinocyte Serum-Free Media; KSFM). Principle component analysis (PCA) depicts that each condition clusters independently from one another and we observe the largest variation between the GBS media control and the GBS adhered to hVECs (**Figure 5B**). Using a fold change cut off ≥ |1.5| and an FDR adjusted p value of < 0.05 as parameters, we identified 201 and 258 genes that were uniquely altered during mono-infection and co-infection, respectively, and 245 genes that were shared (**Figure 5C, Table S1**).

**Figure 4.**
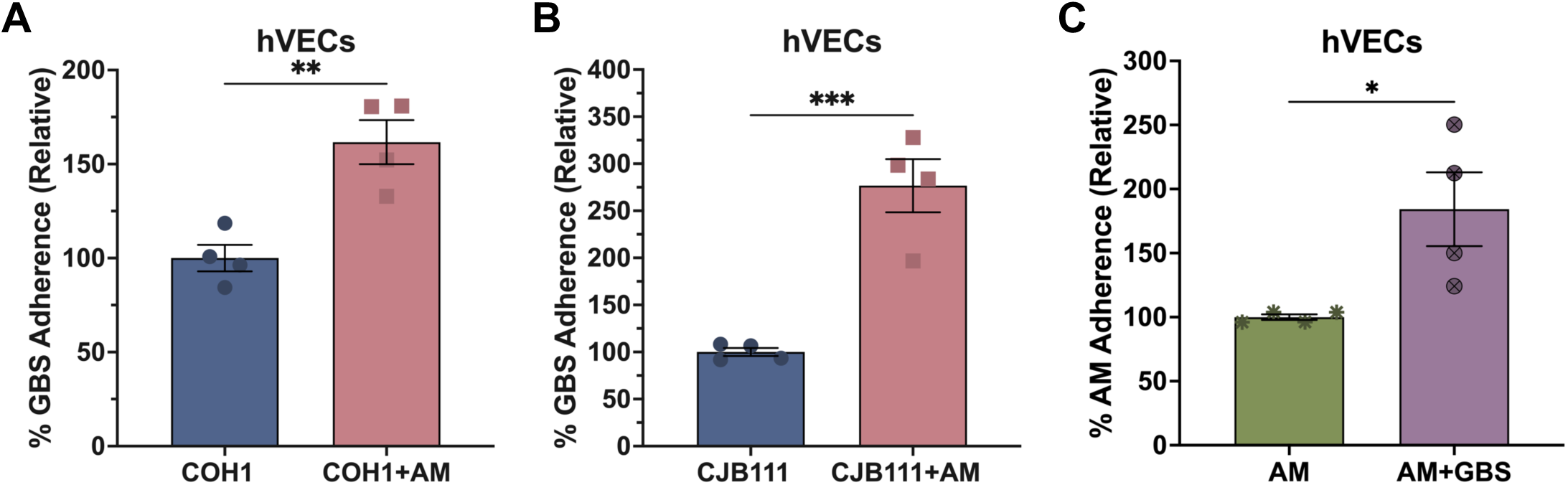
GBS and A. muciniphila synergize on human vaginal epithelial cells (hVECs). Human vaginal epithelial cells were grown to confluency in a 24-well tissue culture plate. GBS and *A. muciniphila* were grown overnight and standardized to an MOI of ∼1, and co-infection was performed at a 1:1 ratio. Adherence for GBS strains **(A)** COH1 and **(B)** CJB111 -/+ *A. muciniphila* or **(C)** *A. muciniphila* -/+ COH1 (normalized to mono-condition) was calculated. Statistical analysis was determined using a Student’s T-test. Data represent the average of independent experiments. Error bars represent +/-SEM. ns, > .05; *, *p* ≤ .05; **, *p* ≤ .01; ***, *p* ≤ .001.

**Figure 5.**
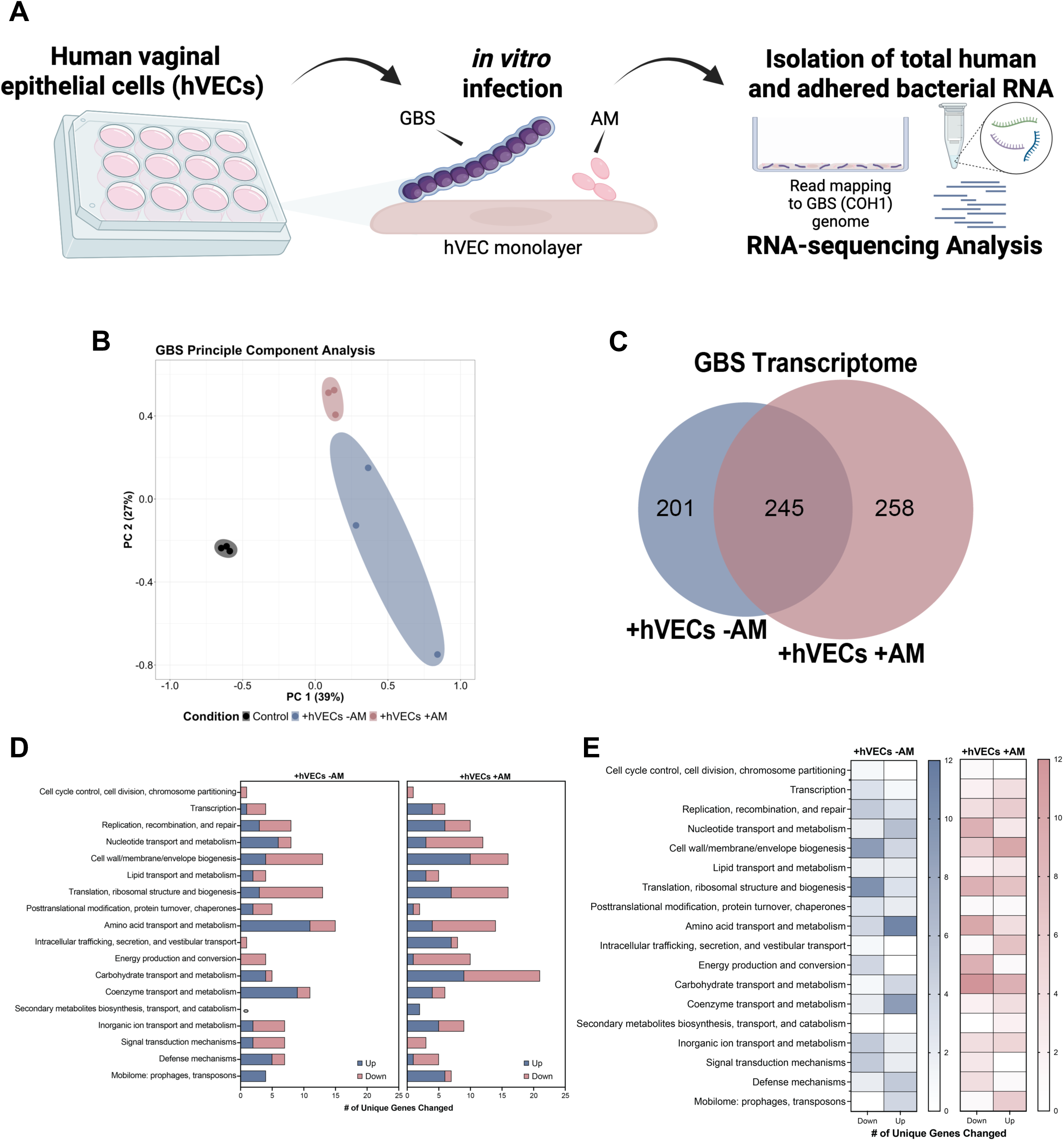
Impact of *A. muciniphila* on the GBS transcriptome. **(A)** 4h mono- and co-infections of human vaginal epithelial cells using GBS (COH1) and *A. muciniphila* at an MOI of ∼100 for RNA-sequencing analysis. **(B)** PCA plot displays gene counts and **(C)** Venn diagram compares transcriptomic changes for GBS under each condition. **(D)** GBS genes unique to each condition with an FDR *p*-value ≤ 0.05 were sorted based on biological pathways and compared to examine the unique effect of *A. muciniphila*. **(E)** Genes unique to each condition with an FDR *p*-value ≤ 0.05 were sorted based on biological pathways and compared to examine the unique effect of *A. muciniphila*. Data represent three biological replicates and differentially expressed genes with an FDR *p*-value ≤ 0.05.

We next organized these uniquely altered genes by orthological groups and broadly compared trends in gene expression between mono-infection and co-infection conditions (**Figure 5D**). Here, we identified significant changes in regulation of carbohydrate transport and metabolism; intracellular trafficking, secretion, and vestibular transport; and energy production and conversion. During mono-infection, genes involved in amino acid transport and metabolism were predominantly up-regulated and shifted to being predominantly down-regulated during co-infection with *A. muciniphila* (**Figure 5E**). Further, we observed a profound shift for genes involved in cell wall/membrane/envelope biogenesis, where the majority of uniquely altered genes belonging to this ontology group were predominantly down-regulated during mono-infection as opposed to being up-regulated during co-infection.

### GBS uniquely modulates cell envelope gene expression in the presence of *A. muciniphila*

We identified numerous genes, both unique to each condition and shared, and examined their expression. We decided to further examine the alterations in expression of select genes involved in cell wall/membrane/envelope biogenesis (**Figure 6A-B**). Here, we noted an interesting difference in regulation of GBS capsule biosynthesis genes, including *cpsA* and *cpsB* which were significantly down-regulated during mono-infection, while *cpsG* and *cpsK* were significantly up-regulated during co-infection. We also observed differences in expression of genes encoding: pilus-island 2b (PI-2b) under both conditions, as well as changes in expression of genes encoding pilus-island 1 (PI-1), plasminogen-binding protein (PbsP), alpha-like surface protein (Rib), serine-rich repeat protein (Srr) 2, and the secretion and glycosylation system for Srr2, which were unique to either condition. Moreover, we found that expression of genes involved in PI-2b synthesis, *bp-2b* and *ap1-2b*, were more up-regulated during co-infection with *A. muciniphila*, as compared to during mono-infection (**Figure 6C**). Conversely, expression of genes involved in PI-1 synthesis, *ap1-1* and *srtC1-1*, were only significantly up-regulated during co-infection.

**Figure 6.**
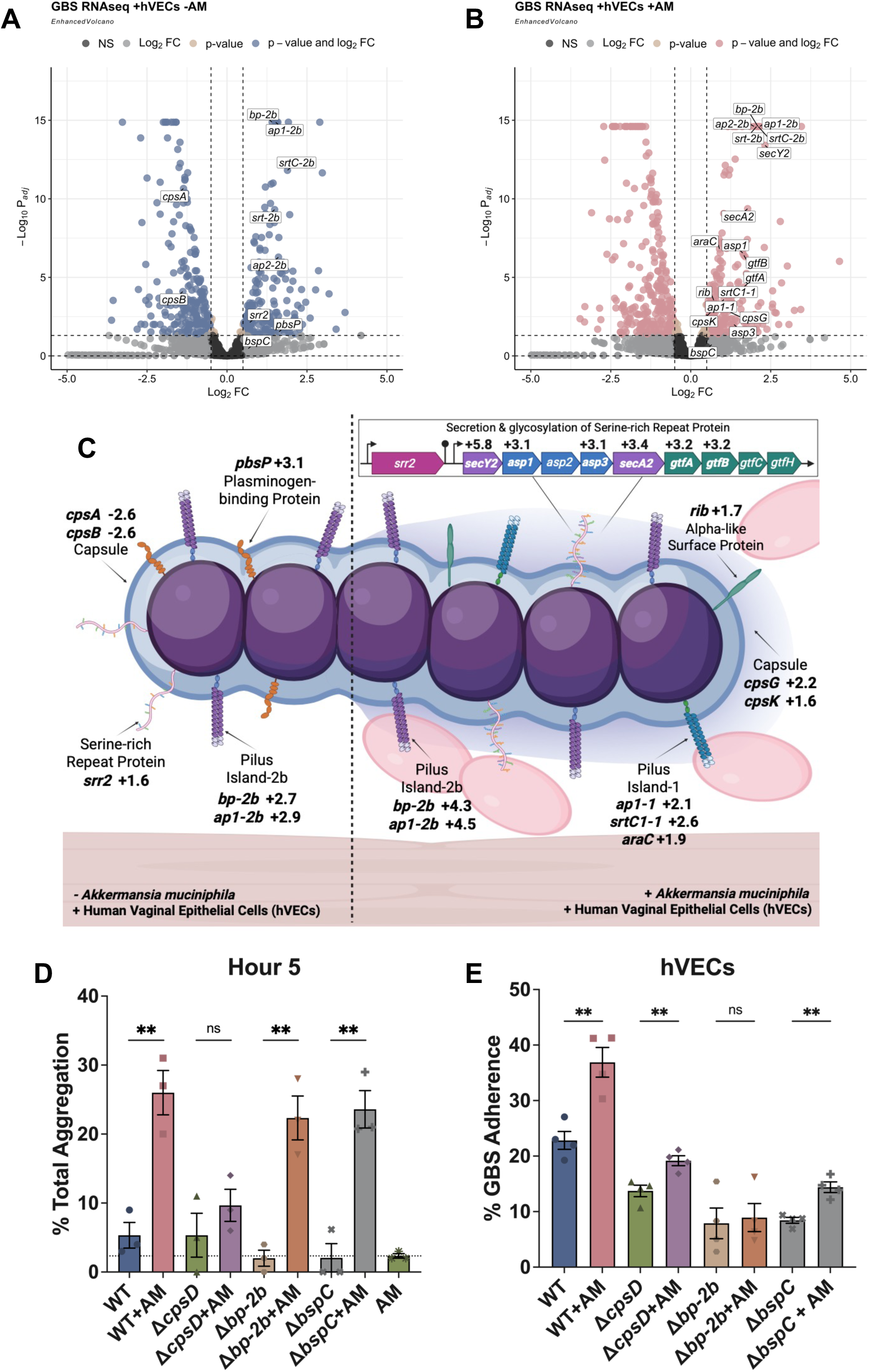
GBS uniquely modulates cell envelope during mono-and co-infection. (A-B) Volcano plots compare the effect of +hVECs –*A. muciniphila* or +hVECs +*A. muciniphila*. **(C)** Select GBS gene fold changes are shown and compared between both conditions. RNAseq data represent 3 biological replicates and differentially expressed genes with an FDR *p*-value ≤ 0.05. Aggregation and adherence was evaluated for various GBS mutants. **(D)** Graph depicts aggregation at 5h for GBS WT and Δ*cpsD* (Hy106), Δ*bp-2b*, Δ*bspC* mutants -/+ *A. muciniphila*. **(E)** Graph depicts percent adherence (raw) for GBS WT and Δ*cpsD* (Hy106), Δ*bp-2b*, Δ*bspC* mutants -/+ *A. muciniphila*. Statistical analysis was determined using a Student’s T-test **(E-F)**. Data points represent the average of independent experiments. Error bars represent +/-SEM. ns, > .05; **, *p* ≤ .01. Panel C was created using Biorender.com.

Another notable difference was in *srr2* expression during mono-infection, as this gene was not significantly altered during co-infection. However, in the presence of *A. muciniphila* we observed the up-regulation of numerous genes involved in the secretion and glycosylation of Srr2, including; *secY2* and *secA2*, which encode subunits for the translocase in this secretion (Sec) system, *asp1* and *asp2*, which encode accessory proteins for this Sec system; and *gtfA* and *gtfB*, which encode a glycosyltransferase and glycosylation chaperone for this Sec system, respectively. Previous work has shown that *srr2* contains a terminator at the end of the coding region, and the downstream Sec system its own promoter^34^. We did not identify differences in notable regulators of virulence, including *saeR*, *saeS* and *covS* **(Table S2)**.

We sought to examine the role of specific GBS representative surface factors, using GBS mutants in capsule (Δ*cpsD*), PI-2b (Δ*bp-2b*), and the Group B streptococcal surface protein BspC (Δ*bspC*), which was used as an example of a gene that was not altered in our RNA-seq analyses, but has previously been shown to contribute to GBS self-aggregation and adherence to hVECs ^33^. We observed that *A. muciniphila* significantly increased GBS aggregation with the Δ*bp-2b* and Δ*bspC* mutant strains like observed with WT GBS but did not significantly increase aggregation with the Δ*cpsD* mutant, suggesting the importance of capsule for this interaction (**Figure 6D**). We further examined the role of these surface factors in adherence to hVECs and observed reduced adherence to hVEC with these mutants as has been published previously^32,33,35^, but that *A. muciniphila* was not able to significantly increase GBS adherence of the Δ*bp-2b* mutant, suggesting the importance of this pilus island to the *A. muciniphila* dependent increase in GBS hVEC attachment (**Figure 6E**).

Thus, we demonstrate that the increases in GBS aggregation and attachment to hVECs in the presence of *A. muciniphila* are dependent on GBS capsule and pili, respectively, and notably *A. muciniphila* induces upregulation of both capsule and PI-2b.

### GBS vaginal colonization is significantly reduced during continual intravaginal treatment with *A. muciniphila*

Our results thus far demonstrate that *A. muciniphila* aggregates with GBS and promotes GBS adherence to vaginal epithelium, which may be modulated by the differential expression of specific GBS factors. Our previous murine model only examined the effect of *A. muciniphila* added prior to or simultaneous to GBS colonization, and due to the recent interest in using *A. muciniphila* as a probiotic therapy^25–27^ we sought to examine the effect of daily *A. muciniphila* vaginal treatment on GBS persistence. Using our well-established murine model of GBS vaginal colonization^36^, CD-1 mice were synchronized to estrus using 17β-estradiol and intravaginally inoculated with GBS or GBS and *A. muciniphila*. Mice were dosed every day with *A. muciniphila* (treatment) or PBS (mock), and the vaginal lumen was lavaged daily to enumerate GBS burden until 5 days post-colonization when FGT tissues were harvested. We found that daily treatment with *A. muciniphila* significantly reduced GBS burdens compared to the mock treatment group and that the decrease is most discernable on Day 5 post-colonization with a 2.2-log reduction in median CFU/ml observed in the *A. muciniphila* treatment group (**Figure 7A**). This corresponded to a significant decrease in GBS colonization over time (**Figure 7B**). Examination of vaginal lavage by fluorescence microscopy showed that GBS appeared to be more aggregated in the presence of *A. muciniphila* (**Figure 7C**). Importantly, the observed decrease in GBS burden with *A. muciniphila* treatment was also observed in vaginal and cervical tissues collected on Day 5 post-colonization with a 2.1-log and 2.2-log decrease in median, respectively (**Figure 7D,E**).

**Figure 7.**
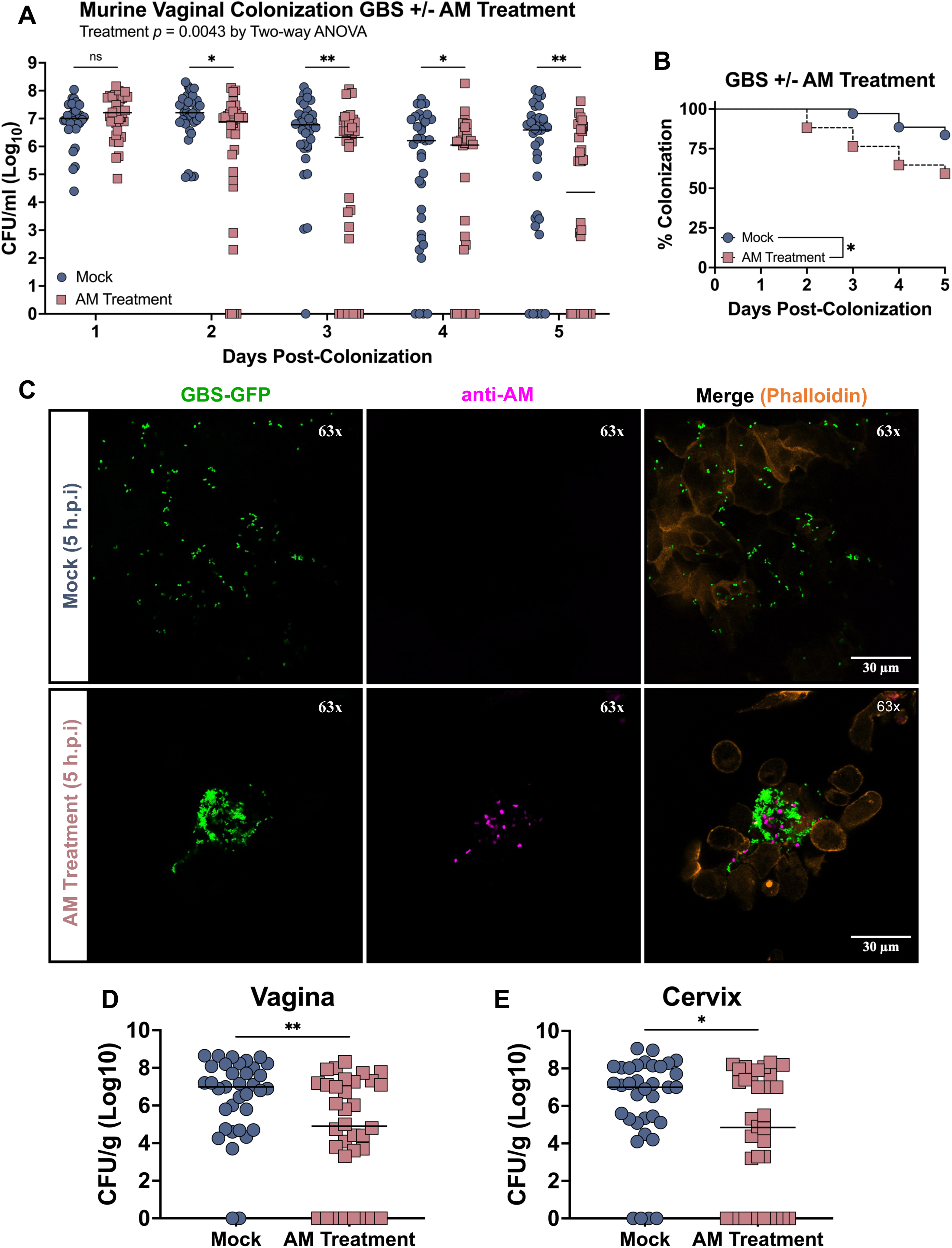
*A. muciniphila* treatment significantly reduces GBS colonization and invasion of the murine cervicovaginal mucosa. **(A)** Recovered GBS CFU from daily lavage of the vaginal lumen and **(B) c**orresponding colonization curve. **(C)** Fluorescence microscopy of pooled vaginal lavage 5h post *A. muciniphila* treatment (representative image). Recovered GBS CFU counts from **(D)** vaginal and **(E)** cervical tissues at day 5. Murine data represent three independent experiments, *n*=10-15 mice per group, per experiment. Microscopy represents 3 biological replicates. Solid lines represent the median. Statistical analysis was performed by two-way RM ANOVA with Uncorrected Fisher’s LSD test **(A)**, Log-Rank test **(C)**, or Mann-Whitney U-test **(D-E)**. ns, > .05; *, *p* ≤ .05; **, *p* ≤ .01.

## DISCUSSION

To date, *A. muciniphila* has been predominantly studied in the context of GI health, specifically for its immunomodulatory role and beneficial effect to human health as a probiotic^25–27^. However, the global effect of *A. muciniphila* in other niches, and its relationship with GBS remains largely understudied. Our lab recently described for the first time the association between *A. muciniphila* and GBS in the vaginal tract, and the co-occurrence of GBS and *A. muciniphila* in pregnant women^22^. Interestingly, work published in 2018 presented the first study that examined any relationship between GBS and *A. muciniphila*, where they showed that an attenuated GBS strain increased the relative abundance of *Akkermansia* from 1% to 28% in the gut of Nile tilapia ^37^. Cross-talk between the gut and genital microbiota of females has been shown to impact the host, and members of the FGT microbiota have been traced back to the rectum which serves as a reservoir ^38^. Notably, research has shown that the same bacterial species have been identified from the rectum and the vagina of pregnant woman, with the vast majority of bacteria demonstrating identical genotypes^39^. As such, it is likely that *A. muciniphila* transiently colonizes the vaginal tract as a result of ‘seeding’ by the gut. Importantly, in addition to our current work identifying co-occurrence of GBS and *A. muciniphila* in pregnant women, recent studies have begun to examine the role of *A. muciniphila* on maternal-fetal health^22^. For example, decreased abundance of *A. muciniphila* has been associated with preeclampsia (PE), a pregnancy specific multisystem disorder that affects 2-8% of pregnancies and is the leading cause of maternal morbidity ^40^. Importantly, it was found that *A. muciniphila* can significantly promote fetal growth and improve placental pathology in a murine model of PE, suggesting the role of probiotic *A. muciniphila* in maternal-fetal health ^40^. This potential new role of *A. muciniphila* in maternal-fetal health led us to further examine its effect on GBS in the FGT..

Collectively, our observations herein further suggests the association between GBS and *A. muciniphila* in the vaginal tract as a very new point of interest. We present additional evidence of GBS and *A. muciniphila* co-occurrence and demonstrate that whilst present in the vaginal microbiota, *A. muciniphila* is not found at a high abundance. This suggests that *A. muciniphila* is likely not a native member of the vaginal microbiota, but a non-native commensal originating from the GI tract. Considering how abundant *A. muciniphila* is in the colon^41^, it is likely that over the course of their lifetime individuals become transiently colonized with *A. muciniphila* in their vaginal tract. Herein, we report for the first time *in vitro* interactions between GBS and *A. muciniphila*.

Our RNA sequencing analysis builds towards understanding the complexities behind GBS colonization and infection of the vaginal tract. We identified important surface factors that were modulated by GBS in the presence of hVECs, as well as nuances in gene expression upon the inclusion of *A. muciniphila*. For example, PI-2b has previously been shown to be necessary for attachment to cells of the FGT, binding to host mucins, and vaginal colonization *in vivo*^42^. We found that PI-2b expression is upregulated to a higher degree during co-infection with *A. muciniphila* compared to mono-infection. It is possible that *A. muciniphila* directly upregulates PI-2b expression, however, it is also possible that *A. muciniphila* modifies the host environment and thereby indirectly upregulates PI-2b expression. Interestingly, it appears that interaction with *A. muciniphila* is dependent on some but not all of the surface factors that we found to be differentially expressed, as PI-2b is required for co-aggregation with *A. muciniphila*, but not for *A. muciniphila-*mediated increase in adherence. These data highlight the complex and contrasting mechanisms that may be involved in interbacterial aggregation as compared to those involved in attachment to host cells.

All of the 10 known capsular serotypes demonstrate distinct molecular compositions that act as a sort of fingerprint. The arrangement and structure of the base monosaccharides (glucose, galactose, and N-acetylglucosamine) vary enough to be antigenically distinct, with the common terminally linked sialic acids (Sias) ^43,44^. We identified numerous GBS genes involved with capsule production to be significantly upregulated during co-infection of hVECs, including a glycosyltransferase involved in adding glucose to the galactose base sugar (*cpsG*) ^43^, and a sialyltransferase that adds the ⍺2,3-linked terminal Sias to the repeating sugar unit (*cpsK*) ^45^. Interestingly, the human GI tract is lined by mucus comprised of heavily O-glycosylated secreted and membrane anchored mucins ^46^, and an increasing gradient of sialylation of *O*-glycans in mucin is observed in the distal colon of humans, a niche that *A. muciniphila* preferentially inhabits ^41,47^. Recent work has shown that the O-glycans on these mucins serve as an attachment point and nutrient source for *A. muciniphila*, and that endogenous sialidase activity of *A. muciniphila* enhances binding to sialylated mucins by removing the Sias. The sialylated capsule protects GBS from complement deposition and phagocytosis, as well as enhances biofilm formation, inhibits the binding of antimicrobial peptides, and alters adherence to cells and mucins^43^. Further investigation is required to determine if *A. muciniphila* cleaves the terminal Sias on GBS capsule, which would likely impact GBS susceptibility to phagocytic clearance. This remains an avenue of our future studies.

As mentioned, live and pasteurized probiotic *A. muciniphila* have become widely popular ^28,29^. We identified that daily *A. muciniphila* vaginal treatment significantly reduces GBS persistence *in vivo*. This decrease in GBS in the vaginal lumen translates to a decrease in invasion of the cervicovaginal mucosa; the vagina and cervix are of particular interest as they are at the forefront of the FGT and are predominantly lined with mucus^48,49^. Further, we observed increased GBS aggregation during *A. muciniphila* treatment *in vivo*, which could be a mechanism for *A. muciniphila* to limit GBS persistence in the vaginal lumen and access to epithelial cells. Our observations provide new evidence for a probiotic role of *A. muciniphila* in the vaginal tract, and potential treatment to limit GBS colonization. Notably, our lab previously demonstrated the probiotic potential of *Streptococcus salivarius* K12, a member of the human oral microbiota which has been isolated from the vaginal tract of pregnant women, albeit in low frequency ^50^.

In sum, we find that *A. muciniphila* has a significant impact on GBS aggregation and colonization in both *in vitro* and *in vivo* conditions. Further studies are required to elucidate the mechanisms by which *A. muciniphila* treatment reduces GBS burden during vaginal colonization, including potential modification of GBS capsule composition through cleavage of Sias as well as modulation of host mucosal immunity. These observations provide a platform for continued studies of *A. muciniphila* in the FGT, which remains a focus for the work in our laboratory.

## MATERIALS AND METHODS

### Bacterial strains and growth conditions

*Streptococcus agalactiae* (GBS) isolates A909 (serotype Ia), H36B (serotype Ib), COH1 (serotype III), NEM316 (serotype III), CJB111 (serotype V), and 41 vaginal isolates from pregnant women were cultured statically in Todd-Hewitt broth (THB; Research Products International) at 37°C throughout this work unless stated otherwise in method. When necessary GBS was grown on Todd-Hewitt agar (THA). *Akkermansia muciniphila* Muc^T^ strain (ATCC BAA-835) was grown in pre-reduced brain-heart infusion (BHI; Research Products International) media supplemented with 0.1% porcine gastric mucin (PGM) at 37°C throughout this work. When necessary *A. muciniphila* was grown on Todd-Hewitt agar (THA). *A. muciniphila* was cultured anaerobically in a Coy Laboratory Products Type A, vinyl anaerobic chamber using an atmospheric gas mix of 85%N_2_/10%CO_2_/5%H_2_ unless stated otherwise in method.

### Tissue culture conditions

Human vaginal epithelial (VK2/E6E7; ATCC Cat# CRL-2616) cell line ^51^were obtained from the American Type Culture Collection and were maintained in keratinocyte serum-free medium (KSFM; Gibco) supplemented with 0.5 ng/ml human recombinant epidermal growth factor and 0.05 mg/ml bovine pituitary extract at 37°C in 5% CO_2_.

### Animal studies

7-week old female CD-1 mice were purchased from Charles River Laboratories. Mice were housed in appropriate ABSL facilities at the University of Colorado Anschutz Medical Campus (CU-AMC) and allowed to acclimate for at least one week prior to experimentation. All animal work was approved by and performed in accordance with the Institutional Animal Care and Use Committee (IACUC) of the CU-AMC under IACUC protocol #00316.

### Metagenomic data analysis

A publicly available vaginal metagenomics data set from pregnant individuals was analyzed for trends between GBS and *A. muciniphila.* Normalized read counts with assigned taxa were downloaded from Baud et al.^31^ and processed to quantify correlative relationships between the presence of different microbes. Samples were binned as positive for a microbe if they contained at least 5 reads assigned to the associated taxon. Number of samples positive for *S. agalactiae*, *A. muciniphila*, both, and neither were used to identify statistical correlation. Normalized read counts for *S. agalactiae* in the *A. muciniphila* positive or negative samples were also assessed.

### Bacterial aggregation assays

GBS and *A. muciniphila* were grown overnight, washed, and standardized to an OD_600_ of 1.0 in phosphate-buffered saline (PBS). Mono-cultures were examined at a 1:1 ratio (Bacteria to PBS) and co-cultures were examined at a 1:1 ratio (GBS to *A. muciniphila*) in 1 ml total volume at room temperature (RT). Samples were mixed vigorously and 10 µl of sample were taken from the top of each sample immediately and diluted in 90 µl of PBS to measure optical density with a plate reader. OD_600_ was examined for hours 1, 3, and 5, and percent aggregation was calculated based on OD_600_ at hour 0.

### Bacterial adherence assays

Bacterial adherence assays were performed to determine the total number of cell-surface adhered GBS as previously described ^35^. GBS was grown overnight and subcultured (1:10) in fresh media to mid-logarithmic phase and standardized to an OD_600_ of 0.4 (CFU∼1x10^9^). *A. muciniphila* was grown overnight and standardized to an OD_600_ of 0.4 (CFU ∼4x10^8^). hVECs were infected at a multiplicity of infection (MOI) of ∼1 and incubated for 30 minutes at 37°C in 5% CO_2_. Co-infection was performed at a 1:1 ratio of GBS and *A. muciniphila*. Following, hVECs were gently washed 4 times with sterile PBS, released from the well with 100 µl 0.25% trypsin-EDTA (Thermo Fisher Scientific), and lysed with 400 µl 0.025% Triton X-100 (Thermo Fisher Scientific) for a total final volume of 500 µl. Cell lysates were serially diluted and plated on THA to enumerate GBS CFU alongside the bacterial input. Percent adherence was calculated based on input.

### Confocal fluorescence microscopy

Samples were examined by fluorescence microscopy to determine spatial distribution of bacterial cells. Bacterial aggregates or murine vaginal lavage samples were fixed in 4% paraformaldehyde for 30 minutes at RT. Samples were washed three times and stored in 1% BSA in PBS until staining. For microscopy, GFP-expressing COH1 and CJB111 strains were used to image GBS in bacterial aggregates and murine vaginal lavage, respectively. For staining, Rabbit anti-*A. muciniphila* polyclonal primary antibody (Sigma-Aldrich) paired with Goat anti-Rabbit IgG Alexa Fluor ™ 633 Secondary Antibody (Sigma-Aldrich) was used to detect *A. muciniphila* and Phalloidin-iFluor 555 (Abcam) to visualize F-actin. Samples were imaged using an LSM 780 inverted laser scanning confocal microscope (Zeiss) with the 63x/NA 1.40 oil immersion objective lens using the 488nm, 561 nm, and 633 nm continuous wave lasers. Image acquisition was performed using Zen Black software and processed using Fiji software.

### Bacterial infection of human vaginal epithelial cells for transcriptomics analysis

GBS was grown in biological triplicate overnight and subcultured (1:10) in fresh media to mid-logarithmic phase and standardized to an OD_600_ of 0.4 (CFU∼1x10^9^). *A. muciniphila* was grown in biological triplicate overnight and standardized to an OD_600_ of 0.4 (CFU ∼4x10^8^). hVECs monolayers were infected at a multiplicity of infection (MOI) of ∼100, plates were centrifuged at 200 x g for 5 minutes and incubated for 4 hours at 37°C in 5% CO_2_. Co-infection was performed at a 1:1 ratio of GBS and *A. muciniphila* (Total bacterial MOI = 200). Input was serially diluted and plated to confirm appropriate MOI. Immediately following infection, supernatant was removed. This was done in order to only retain bacteria actively attached to hVECs. Following, 1 ml of RLT lysis buffer (Qiagen) supplemented with 0.1% BME (Sigma-Aldrich) was added to each well, pipetted up and down vigorously, with light scraping to ensure cell detachment from well, and frozen at -80°C.

### RNA isolation, library preparation, and RNA-sequencing

Frozen samples were thawed on ice and transferred to 2.0-ml conical screw cap tubes with 0.1-mm zirconia beads. Samples were placed into a Mini BeadBeater (BioSpec) and homogenized two times for 40 seconds followed by 1 minute of ice in between each bead-beating step . Samples were centrifuged for 30 seconds at 17,000 x g and supernatant was collected and placed in a new tube containing 95% molecular grade ethanol. Total RNA was prepared using a RNeasy kit (Qiagen) as previously described^52^. Once isolated, DNA was removed using a Turbo DNA-free™ kit (Invitrogen). RNA concentrations were measured using a nanodrop spectrophotometer to ensure a concentration of ≥50ng/µl in 25 µl. RNA samples were sent overnight to SeqCenter (Pittsburg, PA) on dry ice. Samples were treated with Invitrogen DNase (RNAse free), and library preparation was performed using the Stranded Total RNA Prep Ligation with Ribo-Zero Plus kit (Illumina) and 10 bp unique dual indices (UDI). Sequencing was performed using a NovaSeq X Plus, which produced paired end 151 bp reads. Demultiplexing, quality control, and adapter trimming were performed with bcl-convert (v4.1.5). Sequencing statistics were included with raw reads. 50M reads package was utilized for samples containing host cells and 12M reads package was utilized for samples containing only bacterial cells.

RNA-sequencing analysis was performed as previously described ^52^. Raw data (.fastq) files were uploaded to the CLC Genomics Workbench (Qiagen; v21.0.5) for analysis using default settings (mismatch cost, 2; insertion and deletion cost, 3; length and similarity fraction, 0.8). Paired reads corresponding to GBS rRNA, *A. muciniphila* rRNA, and human rRNA were removed by aligning to known rRNA sequences and discarded. The remaining unmapped paired reads (-rRNA) were aligned to the *S. agalactiae* COH1 reference genome (GenBank accession number: NZ_HG939456.1), and expression values were calculated using the RNA-seq Analysis function with default mapping and expression parameters. Experimental comparisons were carried out following quantile normalization by use of the RNA-seq experimental fold change feature. Expression values calculated for each gene are shown as normalized reads per kilobase per million values (RPKM). Differentially expressed genes with an FDR adjusted *p*-value ≤ .05 (determined by CLC Genomics Workbench) and a fold-change ≥ |1.5| were considered for further analysis. Transcripts were annotated using GenBank (Accession: NZ_HG939456.1) and COGs were assigned to differentially expressed genes. PCA plots and volcano plots were generated using R Studio (v 4.3.1.) with packages, *ggplot2*, *ggforce*, *edgeR*, *EnhancedVolcano* . Raw data available through the NCBI GEO repository (Accession: GSE306314).

### Murine vaginal colonization

One day prior to colonization, female CD-1 mice were injected intraperitoneally with 0.5mg β-estradiol in 100μl sesame oil in order to synchronize their estrous cycles and vaginally lavaged with 100μl of sterile PBS by pipetting 50μl of sterile PBS (approximately 8 times up and down) and repeating once more with 50μl of fresh sterile PBS. Undiluted vaginal lavage was spot plated on CHROMagar™ StrepB selective chromogenic media and incubated overnight at 37°C. Any mice positive for GBS following pre-lavage were excluded from the study.

Following synchronization, mice were intravaginally inoculated directly with ∼10^7^ CFU of mid-log phase GBS (CJB111) in 10μl per mouse. For co-inoculation, mice were intravaginally inoculated directly with ∼10^7^ CFU of mid-log phase GBS and ∼10^7^ CFU of stationary phase *A. muciniphila* in 10μl per mouse (5μl per strain). Following inoculation, mice were vaginally lavaged daily with 100μl of sterile PBS by pipetting 50μl of sterile PBS (approximately 8 times up and down) and repeating once more with 50μl of fresh sterile PBS. Vaginal lavage was vortexed briefly, serially diluted 10^-1^ through 10^-4^, track-plated on CHROMagar™ StrepB, and incubated overnight at 37°C. Undiluted samples were spot-plated for every day post-inoculation. Vaginal lavaging and administration of PBS (mock) or *A. muciniphila* (Treatment) was performed daily through experimental end-point. Lavage data represent at minimum two independent experiments with *n*=10-15 mice per group, per experiment.

On day 5 of colonization, the reproductive tract of mice was harvested to enumerate GBS burden. Mice were humanely euthanized by primary CO_2_ and secondary cervical dislocation. Immediately afterwards, vaginal lavage was collected as described above. Following dissection of the female reproductive tract, the vagina, cervix, and uterus were separated and placed into separate 2-ml screw-capped tubes containing ∼1 cm of 1-mm zirconia beads and 500μl sterile PBS. Tubes were weighed before and after adding tissues to calculate tissue weight. Homogenization was performed by bead beating for 1 minute at maximum speed and resting on ice for 1 minute for a total of 2 repetitions. Tissue homogenates were vortexed briefly, serially diluted 10^-1^ through 10^-4^, track-plated on CHROMagar™ StrepB, and incubated overnight at 37°C. Undiluted samples were spot-plated for every tissue. Tissue data are from at least two independent experiments, *n*=10-15 mice per group, per experiment.

### Quantification and statistical analysis

All statistical analysis was performed using Prism software for MacOS (GraphPad Software; v10) and statistical significance was accepted at *p*-values of ≤ 0.05. RNA-seq analysis and statistics were performed on CLC Workbench. Differences with a FDR *p*-value ≤ 0.05 were considered significant in RNA-sequencing analyses, and differences with a *p-*value ≤ 0.05 were considered significant for all other experiments. Number of animals or sample size, bars, p-values, and specific statistical tests are specified in the corresponding figure legends.

## ACKNOWLEDGMENTS

We would like to thank SeqCenter for performing Illumina RNA sequencing; Maddalena Lazzarin, Daniela Rinaudo, and Immaculada Margarit of GSK, Siena, Italy for providing the GBS Δ*bp2b* mutant; Raphael H. Valdivia of Duke University, Durham, NC for his advice and expertise on *A. muciniphila*; and Breck Duerkop of the University of Colorado for facilitating anaerobic culturing. This work was supported by NIH grants R01AI153332 (K.S.D), R21AI186346 (K.S.D), and F32AI186285 (S.M.M).

## SUPPLEMENTAL TITLES AND LEGENDS

**Table S1. All differentially expressed GBS genes with a ≥ |1.5|-fold change and an FDR *p*-value ≤ 0.05. Excel table**

**Table S2. Comparison of select differentially expressed GBS genes in mono- and co-infection conditions with a ≥ |1.5|-fold change and an FDR *p*-value ≤ 0.05.**

